# *Arabidopsis thaliana* interaction with *Ensifer meliloti* can support plant growth under N-deficiency

**DOI:** 10.1101/2020.07.25.221283

**Authors:** Grace Armijo, Tatiana Kraiser, María P. Medina, Diana E. Gras, Ana Zúñiga, Bernardo González, Rodrigo A. Gutiérrez

## Abstract

Nitrogen (N) is an essential macronutrient for plants. Some plant species obtain this nutrient by interacting with N-fixing bacteria. These beneficial interactions are well described in legumes but have also been observed in non-legume plant species that are unable to form root nodules. We studied the expanding role of beneficial plant-bacteria interactions for N-nutrition in the widely used model plant *Arabidopsis thaliana*. We found that the bacteria *Ensifer meliloti* enhanced *A. thaliana* growth under severe N-deficiency conditions, allowing plants to complete their life cycle. Our results showed that bacteria colonize the rhizosphere associated with the epidermis of the plant root. We also demonstrated that *A. thaliana* possesses genes that are critical for this beneficial interaction and are required for plant-growth promotion by *E. meliloti* under N-deficiency.

This work shows association between *A. thaliana* and *E. meliloti* for plant nutrition under severe N-deficiency, and suggests that plants have conserved-molecular mechanisms to interact with N-fixing bacteria to procure N and escape adverse conditions. Under these circumstances, the supply of N via N-fixation is critical for survival, allowing the plant to complete its life cycle. Our findings provide a new framework and an experimental model system that expand our understanding of plant-rhizobia interactions for plant N-nutrition.

## INTRODUCTION

Nitrogen (N) is an essential macronutrient for plant growth and development and one of the main factors limiting plant productivity in natural as well as agricultural systems worldwide (Frink et al., 1999; Hirel et al., 2011). Although N_2_ is abundant in the atmosphere, plants cannot directly access this form of N. Biological N-fixation, an exclusively bacterial trait, is an essential process that transforms atmospheric N_2_ into ammonia, a form of N that is biologically useful for plants (Olivares et al., 2013). N limitation in soils is an essential evolutionary constraint. As a result, plants have developed multiple strategies allowing them to associate with N-fixing bacteria in order to procure better N-nutrition (Boddey et al., 1995; Estrada et al., 2002; Hurek et al., 2002; Iniguez et al., 2004; Kraiser et al., 2011; Pankievicz et al., 2015; Luo et al., 2016; Mus et al., 2016; Van Deynze et al., 2018). These plant-bacteria interactions have been well studied in legumes, which can symbiotically associate with a phylogenetically diverse group of bacteria, collectively called rhizobia, in a species-specific manner (Oldroyd et al., 2011; Wang et al., 2018). Symbiosis is established by the activation of a plant signal-transduction pathway, which includes two families of genes called Nodulating Signaling Pathway (*NSP*) and Nodule Inception (*NIN*). These genes are activated in the presence of nodulation factors secreted by bacteria and when plants grow under N-limiting conditions (Kalo et al., 2005; Smit et al., 2005; Libault et al., 2009). These genes code for transcription factors, regulators of bacterial-infection establishment, and nodule organogenesis (Schauser et al., 1999; Hirsch et al., 2009).

N-fixation in legume plants is a highly specific and efficient process that requires specialized root organs called nodules, where N-fixing bacteria perform N-fixation (Oldroyd et al., 2011). However, numerous examples of other types of associations or interactions have been described in non-legumes, where nodule formation is not required for N-fixation. These interactions range from endophytic to rhizospheric relationships (Bhattacharjee et al., 2008) and are not necessarily restricted to a specific plant compartment. They can occur in organs such as roots, stems, or leaves (Gyaneshwar et al., 2001; Estrada et al., 2002; Iniguez et al., 2004; Caballero-Mellado et al., 2007; Bhattacharjee et al., 2008; Montañez et al., 2009; Pankievicz et al., 2015; Van Deynze et al., 2018). Recently, it was shown that Rhizobiales are consistently present in high relative-abundance and enriched in the root and leaf communities of phylogenetically-diverse plant hosts (Garrido-Oter et al., 2018).

Despite their importance to plant growth, the mechanisms involved in establishing beneficial interactions between non-legume plants and N-fixing bacteria are poorly understood. In this study, we studied the role of beneficial plant-bacteria interactions for N-nutrition in the widely-used model plant *Arabidopsis thaliana*.

## RESULTS

### Metabolically active *E. meliloti* can promote plant growth in the absence of a N source in the media

In an effort to understand the importance of N-fixation for non-legume plant growth, we evaluated the possibility that the widely used model plant *A. thaliana* could establish beneficial interactions for N-nutrition with N-fixing bacteria. In order to evaluate this possibility, we selected five different bacterial species known to fix N in association with plants and assessed their effect on *A. thaliana* plant-growth under N-limiting conditions: *Ensifer meliloti* RMP110 (Yuan et al., 2006), *Rhizobium etli* CFN42 (Poupot et al., 1995), *Cupriavidus taiwanensis* LMG 19424 (Marchetti et al., 2011), *Paraburkholderia xenovorans* LB400 (Sawana et al., 2014) and *Paraburkholderia vietnamiensis* G4 (Sawana et al., 2014). Two bacteria unable to carry out biological N-fixation were also tested as controls: *Paraburkholderia-phytofirmans* PsJN, known to enhance Arabidopsis growth under standard conditions with full nutrition (Zuniga et al., 2013; Sawana et al., 2014) and *Cupriavidus pinatubonensis* JMP134, capable of associating with plants but without any positive impact on plant growth (Ledger et al., 2012; Zúñiga et al., 2018).

Plants were grown on Murashige and Skoog (MS) media for seven days with 5 mM of KNO_3_ as the only N source to ensure that the seedlings develop properly before treatment, and depend exclusively on media nutrients. The plants were then transferred to MS media without N (MS-N), MS-N supplemented with 2,5mM NH_4_NO_3_ or MS-N inoculated with different bacterial strains. Biomass as the plant dry weight was evaluated seven days after transfer. As seen in Figure 1A, plant dry weight was significantly higher in the presence of *E. meliloti* as compared to non-inoculated media under N-limiting conditions (p < 0.01). Moreover, plant biomass was comparable to that achieved in media supplemented with NH_4_NO_3_ under the same experimental conditions. None of the other bacterial species tested, nor heat-killed *E. meliloti*, increased *A. thaliana* biomass, measured as dry weight (Figure 1A, B). *A. thaliana* plants can grow, flower, and even set viable seeds in the absence of N when inoculated with *E. meliloti* (Figure 1C and Supplemental Figure 1A). Thus, these results show that the enhancement of Arabidopsis plant-growth in the absence of N requires a metabolically active *E. meliloti* and is not caused by the mere presence of neutral or plant growth-promoting bacteria or other N-fixing bacteria.

**Figure 1:**
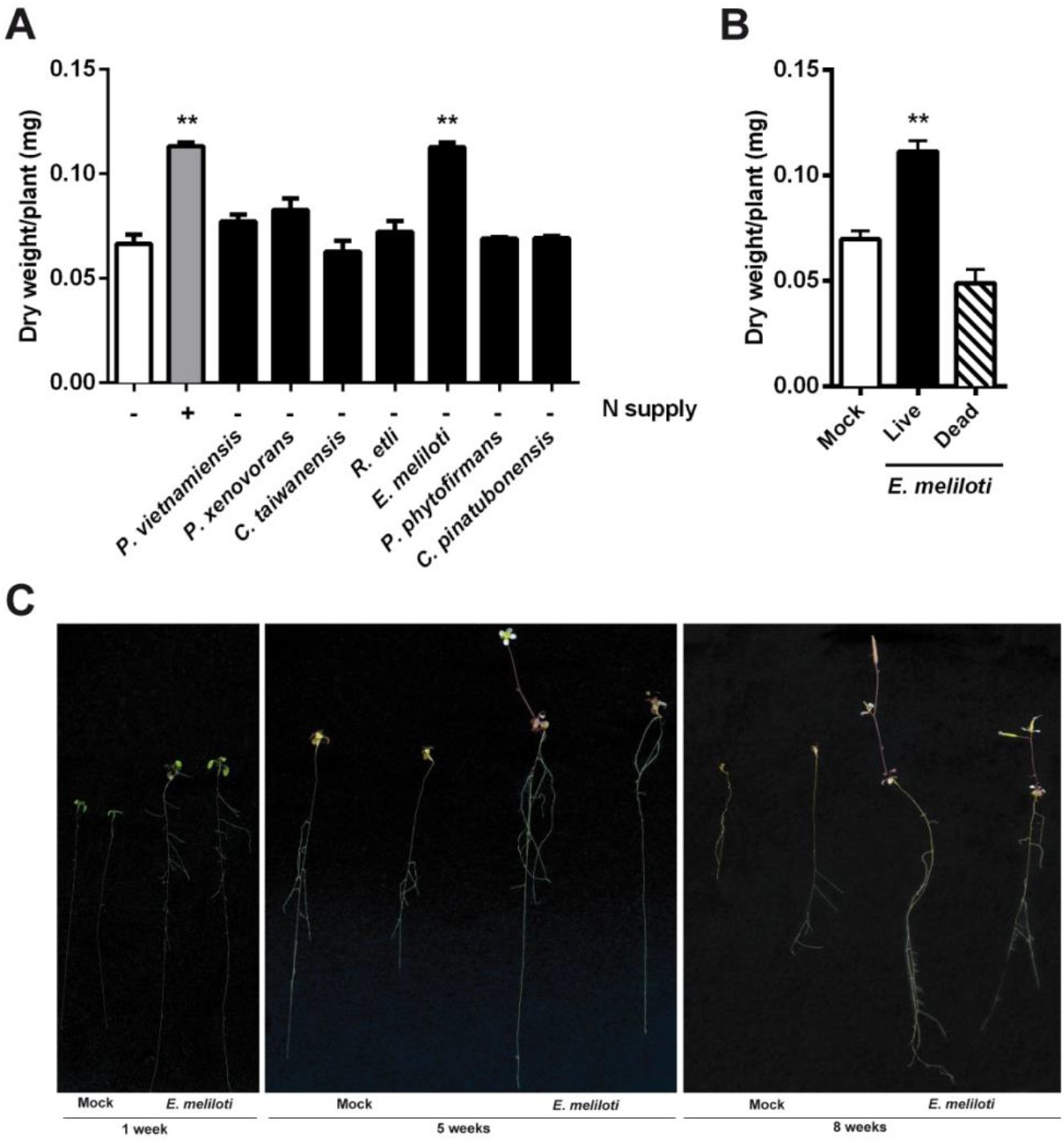
Metabolically active *E. meliloti* can promote plant growth in the absence of a N source in the media. **(A)** One week seedlings grown in KNO_3_ 5mM were transplanted to MS without N, supplemented with 2.5mM NH_4_NO_3_, and to MS without N, inoculated with different bacterial strains (N-fixing bacteria: *P. vietnamiensis* G4, *P. xenovorans* LB400, *C. taiwanensis* LMG19424, *R. etli* CFN42, *E. meliloti* RMP110; no N-fixing bacteria: *P. phytofirmans* PSJN, *C. pinatubonensis* JMP134). Biomass was measured as dry weight seven days after transferring the plants. **(B)** Dry weight of plants inoculated with viable or dead cells of *E. meliloti* or mock-inoculated under the experimental conditions described in (a). **(C)** The effect of *E. meliloti* on the promotion of plant growth without N as compared to mock-inoculated plants at 1, 5, and 8 weeks after transfer. Values plotted correspond to the mean of three independent biological replicates ± standard error. Results were subjected to one-way analysis of variance (ANOVA) and Tukey’s multiple comparison test. The asterisks indicate that the means differed significantly compared to non-inoculated plants grown without N (** p < 0.01). Signs - and + represent N supply.

### Nitrogen fixation enhances *A. thaliana* growth under N-limiting conditions

To determine whether N-fixation was needed for plant-growth promotion under N-limiting conditions, we generate an *E. meliloti nifH* mutant strain via homologous integration of a suicidal plasmid in the bacterial DNA. By disrupting the structural component of nitrogenase, NifH nitrogenase Fe protein, *NifH* gene, we aimed to disrupt the functionality of the entire cluster (**Supplemental Figure 1B**). PCR amplification and direct sequencing were carried out to verify plasmid integration and gene interruption **(Supplemental Figure 1C)**. Alternatively, this strain was tested in symbiosis with alfalfa to confirm *nifH* mutation and lack of N-fixation capacity (**Supplemental Figure 1E)**. As expected, *nifH* interruption was confirmed, and the alfalfa phenotype showed senescent nodules as previously reported (Hirsch et al., 1983). Bacterial viability was also evaluated through growth curves (**Supplemental Figure 1D)**, reaching the same level of growth as the WT strain at 72 hours.

Later, *A. thaliana* plants were inoculated in MS-N with *E. meliloti nifH* mutant and WT strains, and plant biomass was evaluated as described above. As shown in **Figure 2A**, *E. meliloti nifH* had a significantly reduced impact on plant growth as compared to WT strain under N-limiting conditions (p < 0,001), although it still retained some growth-promoting properties. Both WT and *nifH* mutant bacterial strains increased lateral root density compared with mock-inoculated plants. This result suggests the existence of additional bacterial growth-promoting activity that is independent of N-fixation **(Figure 2B)**. To evaluate this possibility, alfalfa, and Arabidopsis plants were inoculated with *E. meliloti* in the presence of N. As expected in alfalfa, nodulation was inhibited (Streeter and Wong, 1988), but plant growth was higher when the bacteria were present **(Supplemental Figure 2).** The same effect on growth was observed in Arabidopsis plants **(Supplemental Figure 2)**.

**Figure 2:**
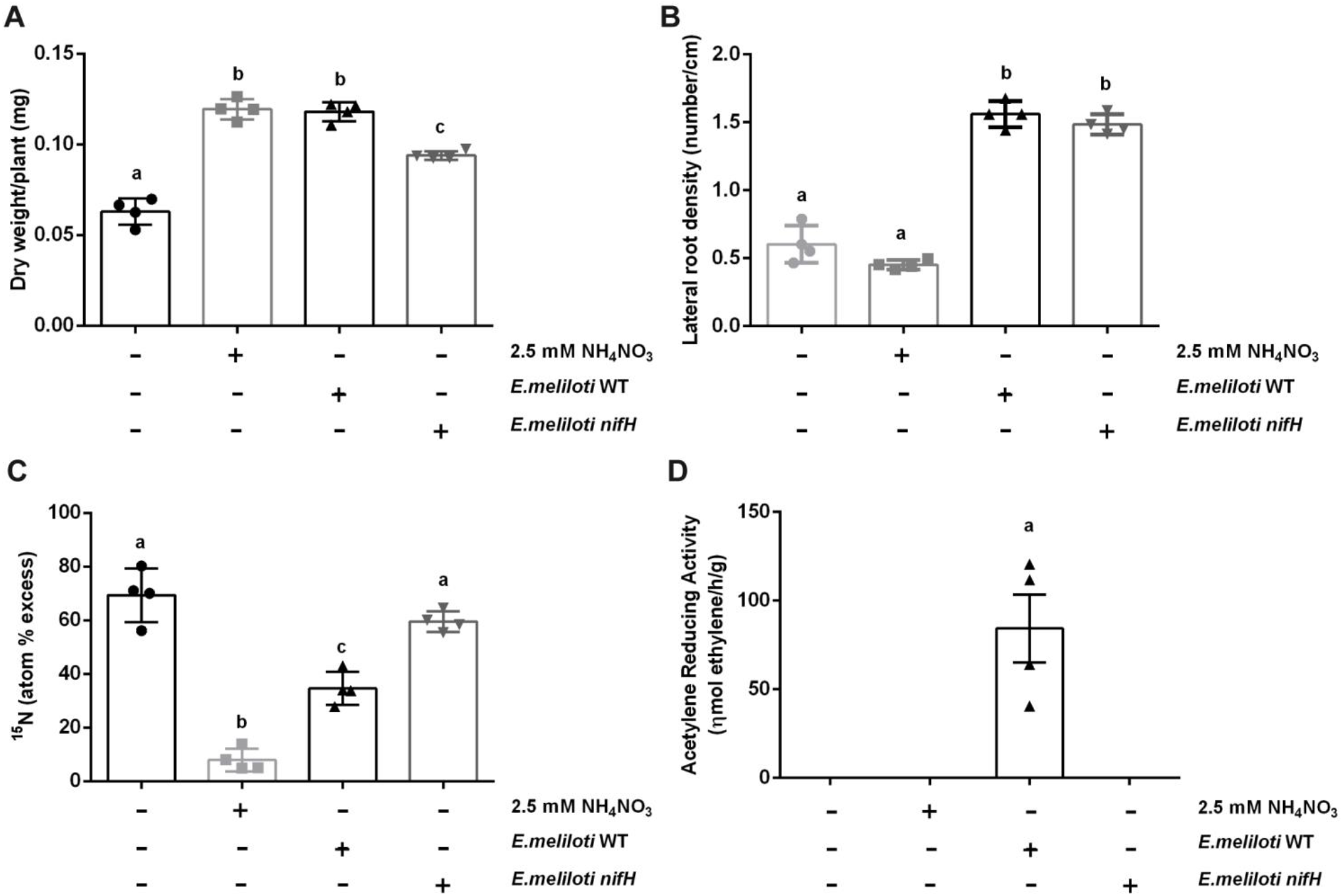
Nitrogen fixation is required for maximal enhancing of plant growth under N-limiting conditions. **(A)** One-week seedlings grown in KNO_3_ 5mM were transplanted to MS without N, supplemented with 2.5mM NH_4_NO_3_, and to MS without N inoculated with WT or *nifH* mutant *E. meliloti* strains. Seven days after transferring the plants, biomass was measured as dry weight and normalized by the number of plants for each condition. **(B)** The primary-root length and number of lateral roots were determined to analyze lateral-root density in the same conditions described above. **(C)** ^15^N isotope dilution assay to estimate biological N-fixation. Plants were grown in KNO_3_ (enriched with 5% of ^15^N) as the only N source and transferred to MS plates without N, supplemented with 2.5mM NH_4_NO_3_, and to MS without N inoculated with WT or *nifH* mutant *E. meliloti* strains. Seven days after transfer, ^15^N isotopic composition in plant tissues was determined using mass spectrometry. Atom percent excess was calculated by subtracting the mean atom% value of the control plants from the atom% of the labeled plants. **(D)** Acetylene Reduction Assay (ARA) as an indirect method to evaluate nitrogenase activity was performed in closed glass-bottles under the same conditions described in (a). Average acetylene reduction rates (ηmol ethylene /h/g) were determined for each treatment. Values plotted correspond to the mean of independent biological replicates ± standard error (biological replicates were graphed, for better analysis). Results were subjected to one-way analysis of variance (ANOVA) and Tukey’s multiple comparison test. The letters indicate a statistical difference between treatments. The p values for each panel are as follows: **(A)** p < 0.001; **(B)** p < 0.001; **(C)** p < 0.01 and **(D)** p < 0.05. Signs - and + represent N supply and presence of the WT and *nifH* mutant bacteria.

To assess whether *E. meliloti* provides fixed N to *A. thaliana* under N-limiting conditions, we performed a ^15^N-dilution assay. Plants were grown on MS-N supplemented with 5 mM of isotopically labeled K^15^NO_3_ (5% ^15^N) as the only N source. They were then transferred to MS media without N and inoculated with WT and *nifH E. meliloti* strains. ^15^N isotopic composition was determined in plant tissues using mass spectrometry seven days after transferring the plants. Higher N incorporation by the plant implies a higher ^15^N dilution. As shown in Figure 2C, and as expected, the highest dilution of ^15^N was observed when plants were transferred to 2.5 mM ^14^NH_4_^14^NO_3_ (sufficient N condition) due to the incorporation of ^14^N readily available in the media. However, in the presence of WT *E. meliloti*, plants showed a reduced ^15^N isotopic proportion as compared to *nifH* and non-inoculated plants (p > 0,01).

Bacterial cells used for inoculation were not isotopically labeled. Therefore, we evaluated as a control, whether the total N contained in the inoculated *E. meliloti* could explain, at least in part, the isotopic dilution observed in *A. thaliana*. Total N in the *E. meliloti* biomass used for inoculation was quantified by mass spectrometry, and this value was used for a theoretical estimate of ^15^N dilution (using the natural abundance of 0.3663% ^15^N (Mariotti, 1983)) under the premise that *A. thaliana* incorporates 100% of the N contained in bacterial cells. As shown in **Supplemental Figure 3**, the total N in *E. meliloti* biomass could not significantly dilute the isotopic composition of *A. thaliana* under these experimental conditions.

As an indirect method to evaluate nitrogenase activity under our experimental conditions, the acetylene reduction assay (ARA) (Hardy et al., 1968) was performed in the same conditions described above. After two weeks of incubation, treated samples and controls were injected with acetylene, and after 24 hours, the concentration of ethylene generated was determined by gas chromatography.

Because the nitrogenase complex can catalyze the reduction of acetylene to ethylene, the amount of ethylene produced is used as an indirect measure of N-fixing activity (Danso, 1995). As shown in **Figure 2D**, we detected acetylene reduction activity when *A. thaliana* was inoculated with WT *E. meliloti* but not with the *nifH* mutant *E. meliloti* strain (p < 0.05).

These results indicate that *A. thaliana* can functionally interact with *E. meliloti* to enhance plant growth under N-limiting conditions via N-fixation. This functional interaction contributes to increased Arabidopsis plant biomass.

### *E. meliloti* colonizes *A. thaliana* root surface

In order to understand the nature of the association between *E. meliloti* and *A. thaliana*, bacterial localization was evaluated by microscopy. Inoculated plants were fixed in glutaraldehyde and evaluated using optical and electron microscopy **(Figure 3A, B, and C)**. For optical microscopy, root longitudinal cuts were made and stained with toluidine blue. Bacterial colonization in the outer layers of the root, including in root mucilage, was observed **(Figure 3A)**. Transmission electron microscopy confirmed the presence of bacterial cells on the surface of plant roots **(Figure 3B)**. *E. meliloti* colonization using scanning electron microscopy was also evaluated, and bacterial cells were preferentially located at the junctions between two epidermal cells of the plant root, in abundant as well as in small groups **(Figure 3C)**. Finally, these results were confirmed by confocal laser scanning microscopy, using a bacterial-strain constitutively expressing Green Fluorescent Protein (GFP), *E. meliloti* P_*lac*Iq_::*GFP*. Consistent with optical and electron microscopy results, no bacterial presence was detected inside the roots, but a significant bacterial presence was detected in the rhizoplane, particularly between epidermal cells **(Figure 3D)**. These results strongly suggest that *E. meliloti* localizes in the *A. thaliana* rhizosphere.

**Figure 3:**
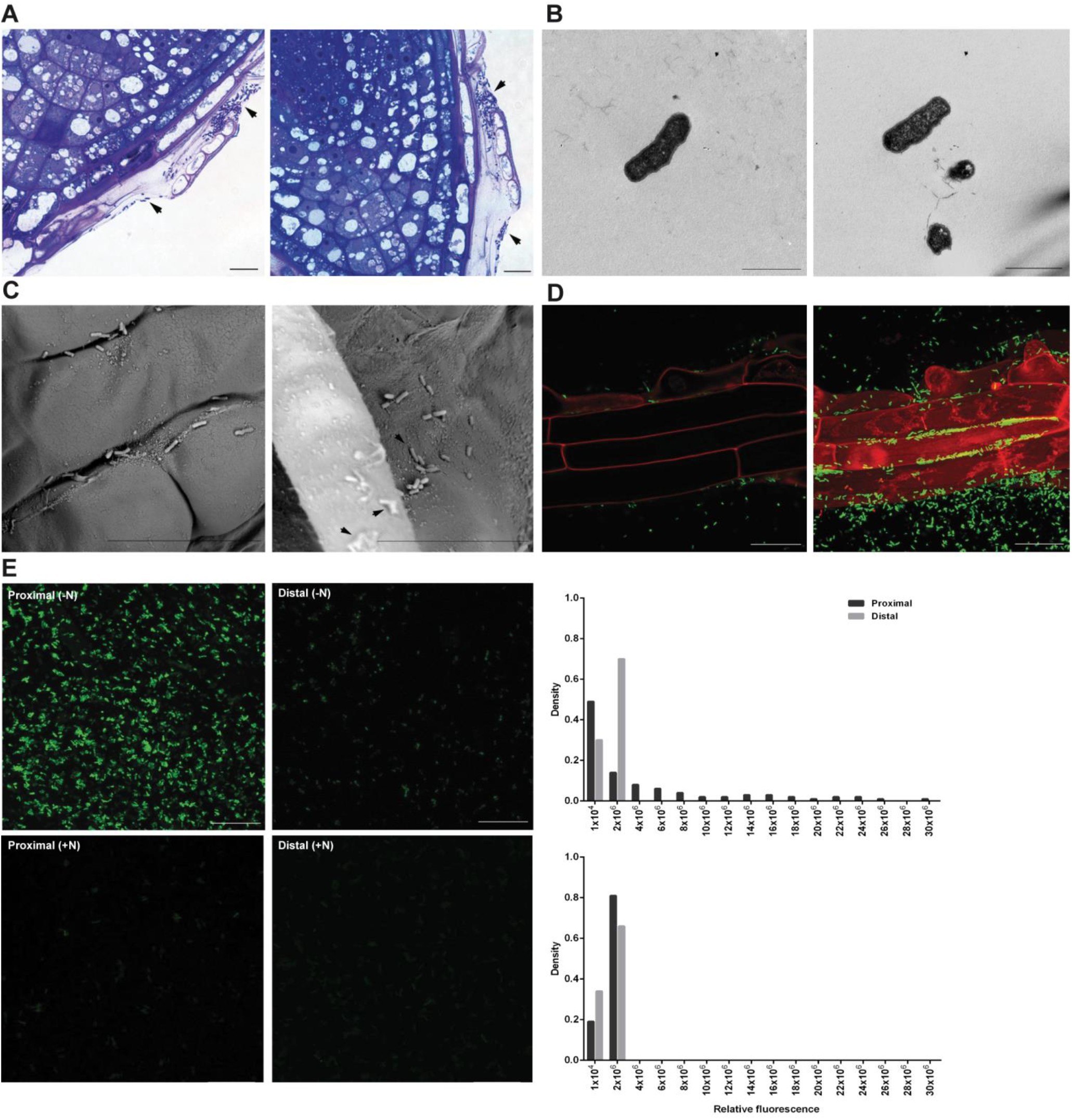
*E. meliloti* colonizes the root surface of *A. thaliana*. Seven-day plants were transferred to MS media without N and inoculated with wild type E. *meliloti* **(A, B** and **C)** or the same strain constitutively expressing GFP (P_*lac*Iq_::*GFP*) **(D)** or expressing GFP under the control of the *Nif*H I promoter (P_*Nif*H_::*GFP*) **(E)**. Two weeks after transferring, *Arabidopsis* roots were fixed in glutaraldehyde and stained with toluidine blue for optical microscopy **(A)**, analyzed by transmission electron microscopy **(B)** or scanning electron microscopy **(C).** GFP-expressing bacteria were visualized using confocal laser scanning microscopy **(D** and **E)**, and epidermal root cells were stained with propidium iodide **(D)**. In **(E)** analysis of agar samples adjacent to *A. thaliana* roots (proximal, left panels) and at a distance of 5 cm (distal, right panels) away from plant roots, was performed in the absence (-N) or presence of N (+N). Z-stacks are showed. Sample fluorescence was quantified and normalized by bacterial area. The frequency density versus relative fluorescence was plotted for −N condition (top graph) and +N condition (bottom graph). Different slides are shown in left and right panels for **(A)**, **(B),** and **(C);** bar represents 10 μm, 1 μm and 20 μm, respectively. In **(D),** a single slide and a z-stack are shown; the bar represents 10 μm. In **(E),** z-stacks are shown; the bar represents 10 μm. Arrows indicate bacterial structures.

To assess whether expression of the *E. meliloti Nif* operon is active on the *A. thaliana* rhizosphere, an *E. meliloti* strain that expresses GFP under the control of the *NifH* promoter (P_*nif*H_::*GFP*) was also generated. *A. thaliana* plants were inoculated with this bacterial derivative as described above, and confocal laser-scanning microscopy analysis of different sites on the agar plates - adjacent to *A. thaliana* roots (proximal) or at a distance of 5 cm from the roots (distal) - were performed **(Figure 3E)**. Differential expression of GFP fluorescence was observed when comparing proximal and distal sites. As a control, the same experiment was made in the presence of N (2.5 mM NH_4_NO_3_), taking sites on the agar plates that were adjacent or distal to *A. thaliana* roots **(Figure 3E)**. For each agar sample, Z-axis slides were taken and integrated with the ImageJ FIJI program (Schneider et al., 2012). A fluorescence threshold was defined, quantified, and normalized by the bacterial area. Then, the frequency and density versus relative fluorescence was plotted for each condition **(Figure 3E)**. We observed a distribution of bacteria with higher fluorescence in agar samples proximal to the plant roots in the absence of N, as compared to the agar samples distant to the plant root or the N-sufficient condition. This suggests that proximal to the plant root, there is an increased nitrogenase activity.

These results indicate that, under our experimental conditions, *E. meliloti* colonizes the root surface of *A. thaliana*. In addition, proximity to the plant root induces *nifH* expression in neighboring bacteria in the absence of N, and such *nifH* gene expression levels significantly decrease away from the plant root.

### *E. meliloti* NodA factor is not required for enhancing *A. thaliana* growth under N-limiting conditions

*NodABC* genes encode proteins responsible for synthesizing core Nod factors. NodA is a host-specific determinant of the transfer of fatty acids in Nod factor biosynthesis and is essential for nodulation (Ritsema et al., 1996). To determine if *Nod* genes are needed for plant-growth promotion under N-limiting conditions, an *E. meliloti nod*A mutant strain was generated via homologous integration of a suicidal plasmid. By disrupting the *Nod*A gene, the first gene in the Nod cluster coding for the acyltransferase NodA, we aimed to disrupt the functionality of the entire cluster and thus to impair the strain’s ability to synthesize Nod signal molecules (**Supplemental Figure 1B**). PCR amplification and direct sequencing were carried out to verify plasmid integration and gene interruption **(Supplemental Figure 1C),** as described above. Also, bacterial viability was evaluated by plotting growth curves (**Supplemental Figure 1D)**. Alternatively, this strain was tested in symbiosis with alfalfa to confirm *nodA* mutation and its inability to form nodules (**Supplemental Figure 1E)**. Subsequently, *A. thaliana* plants were inoculated with *E. meliloti* WT, and *nodA* mutant strains and plant dry weight was evaluated as described above. As shown in **Figure 4**, *nodA* mutant strains can promote plant-growth under N-limiting conditions similar to what we observed with *E. meliloti* WT.

**Figure 4:**
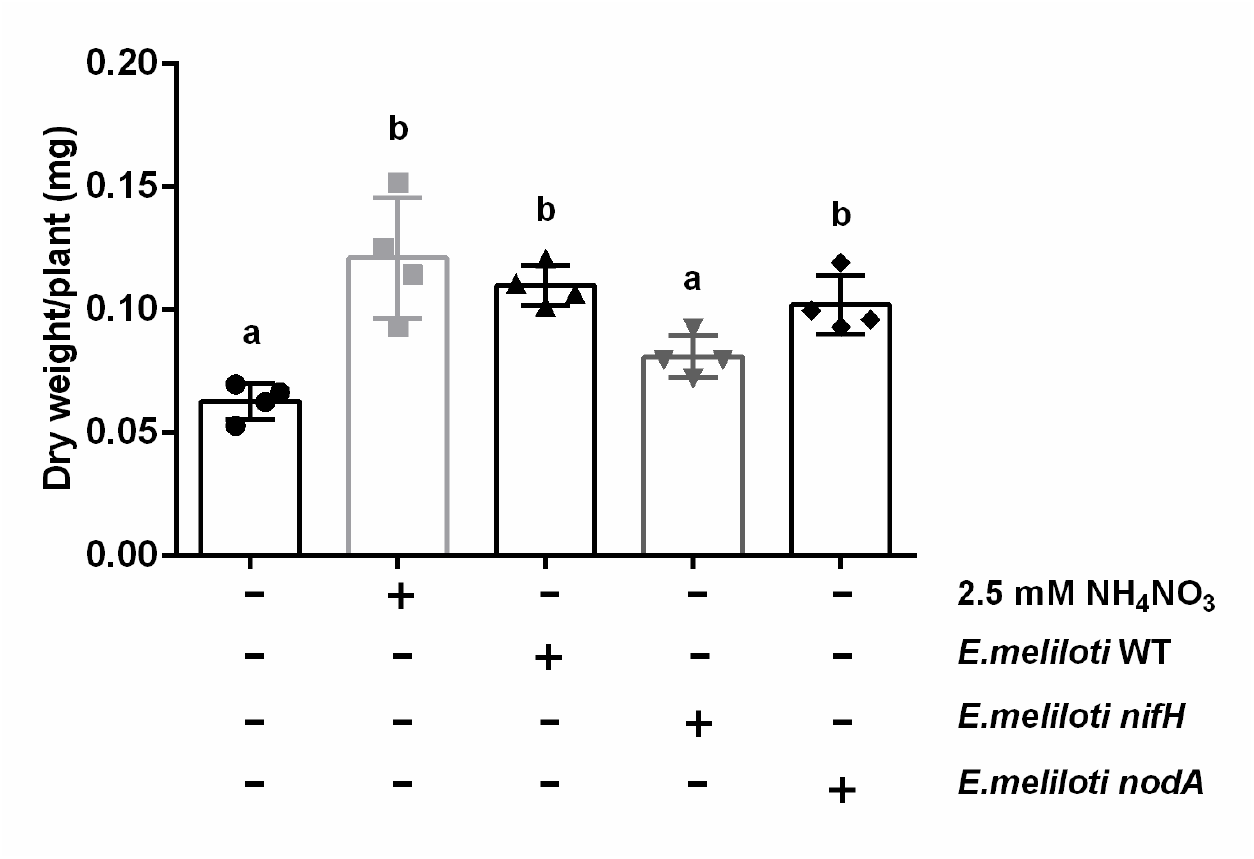
*E. meliloti* NodA factor is not required for enhancing *A. thaliana* growth under N-limiting conditions. One-week seedlings grown in KNO_3_ 5mM were transplanted to MS without N, supplemented with 2.5mM NH_4_NO_3_, and to MS without N inoculated with WT, *nifH*, or *nodA* mutant *E. meliloti* strains. Seven days after transfer, plants were harvested, dried, and weighed to determine biomass. Values plotted correspond to the mean of independent biological replicates ± standard error. Results were subjected to one-way analysis of variance (ANOVA) and Tukey’s multiple comparison test. The letters indicate a statistical difference between treatments (p < 0.05). Signs - and + represent N supply and presence of the WT and *nifH* and *nodA* mutant bacteria.

These results indicate that, under our experimental conditions, *E. meliloti* colonization of *A. thaliana* roots does not depend on the recognition of Nod factors, which are required for bacterial colonization in legumes.

### Arabidopsis *NSP* and *NIN* orthologs of legumes genes are induced upon bacterial inoculation

N-fixation in legume species depends on sophisticated molecular mechanisms that control when and how the symbiotic association is established with rhizobia. Although Arabidopsis plants cannot make nodules, the *A. thaliana* genome contains a set of genes related to those found in legume species, including Nodulation-Signaling Pathway (NSPs) (Delaux et al., 2014) and NIN-like protein (NLPs) genes (Schauser et al., 2005). We conducted a phylogenetic analysis of NSPs and NLPs homologs in *A. thaliana. AtNSP1* (At3g13840) and *MtNSP1* form a subfamily in the NSP tree **(Supplemental Figure 4A)**, supported with bootstrap values of 98% (maximum likelihood), which is consistent with previously published data (Smit et al., 2005; Liu et al., 2011). A similar result was obtained for *AtNSP2* (At4g08250), which forms a separate clade from NSP1 with *MtNSP2a* and *MtNSP2b* **(Supplemental Figure 4A)**. In legumes, NSP1 and NSP2 regulate expression of *NIN* genes. Interestingly, the *A. thaliana* genome encodes nine *NIN*-like protein (*NLPs*) genes (Schauser et al., 2005).

*NLP* and *NIN* genes are part of a conserved subfamily of transcription factors (Schauser et al., 2005). *AtNLP9* and *AtNLP8* fall within the same clade as *MtNLP1 and MtNLP2*, supported with bootstrap values of 98% (maximum likelihood) **(Supplemental Figure 4B)**. *MtNLP1 and MtNLP2* are *AtNLP9* orthologs with 52 and 50% identity, respectively. *AtNLP6* and *AtNLP7* form a separate clade with *MtNLP3*.

To analyze the function of putative *AtNSPs* and *AtNLPs* homologs, we evaluated the expression of these genes in plants transferred to 2.5 mM NH_4_NO_3_ or MS-N medium in the presence or absence of *E. meliloti* WT and harvested 3 and 7 days after the transfer **(Figure 5)**. Results show that *AtNSP1* gene expression is induced when plants are transferred to MS-N medium inoculated with *E. meliloti* WT but not under other experimental conditions **(Figure 5A)**. In contrast, *AtNSP2* was not regulated under the experimental conditions tested **(Figure 5B)**. We chose to analyze expression of the closest homologs to *NIN* genes *AtNLP1* (At2g17150), *AtNLP2* (At4g35270), *AtNLP3* (At4g38340), *AtNLP4* (At1g20640), *AtNLP5* (At1g76350), *AtNLP8* (At2g43500) and *AtNLP9* (At3g59580) (Schauser et al., 2005) under the same experimental conditions described above **(Figure 5 C-I)**. We decided not to test *AtNLP6* nor *AtNLP7* because both genes have been shown to play a key role in nitrate-signaling in *A. thaliana*, accumulating earlier in the nucleus and regulating nitrate-inducible gene expression (Marchive et al., 2013; Guan et al., 2017). We wanted to evaluate homologous genes that were involved in the interaction with the bacteria.

**Figure 5:**
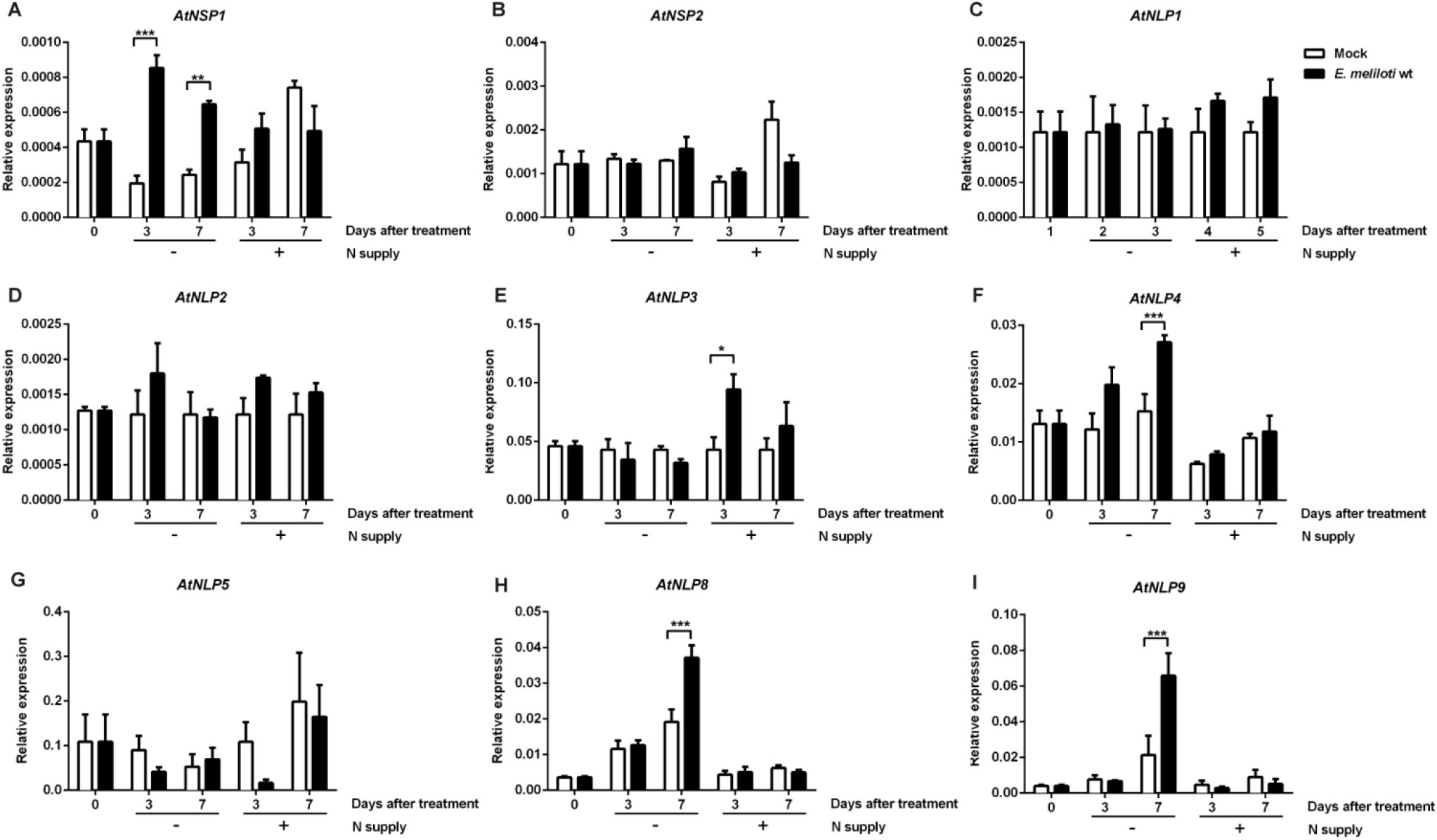
*A. thaliana Nodulating Signaling Pathway (NSP*) and *Nodule Inception-like Protein* (NLP) orthologs of legumes genes are induced upon bacterial inoculation. Plants were grown under sufficient (2.5 mM NH_4_NO_3_) o limiting N conditions and inoculated or not with *E. meliloti* WT. Gene expression was measured using real-time quantitative reverse transcription PCR (qRT-PCR) three and seven days after transferring the plants to the experimental conditions indicated. Treatments were performed as reported previously. Symbols - and + represent N supply. Values plotted correspond to the mean of three independent biological replicates ± standard error. Results were subjected to two-way analysis of variance (ANOVA) and Tukey’s multiple comparison test. The asterisks indicate means that differed significantly as compared to non-inoculated plants (*p < 0.05; ** p < 0.01; ***p<0.001).

Results show gene expression of *AtNLP4, AtNLP8* and *AtNLP9* are induced under N-limiting conditions in the presence of *E. meliloti* **(Figure 5F, H** and **I**, respectively) similar to *AtNSP1* **(Figure 5A)**. In contrast, *AtNLP3* was induced under N-sufficient conditions but only in the presence of bacteria **(Figure 5E)**. Gene expression of *AtNLP1, AtNLP2*, and *AtNLP5* did not change significantly under the experimental conditions evaluated (**Figure 5C, D,** and **G**, respectively). These results suggest a possible function for *AtNSP1*, *AtNLP4*, *AtNLP8*, and *AtNLP9* in *A. thaliana-E. meliloti* interaction under N-limiting conditions.

### Putative transcription factors of *A. thaliana* are essential for functional interaction with *E. meliloti*

To address the possible function of *AtNSP1*, *AtNLP4*, *AtNLP8*, and *AtNLP9* genes in the context of *A. thaliana-E. meliloti* interaction, homozygous mutant lines of *A. thaliana* for *AtNSP1* (salk_036071C (*nsp1.1*); salk_023595C (*nsp1.2*)), *AtNLP4* (salk_100786C (*nlp4.1*); salk_063595C (*nlp4.2)), AtNLP8* (salk_031064C (*nlp8)*) and *AtNLP9* (salk_025839C (*nlp9.1*); salk_042082C (*nlp9.2*)) genes were inoculated with *E. meliloti*, as previously described.

Results show plant-growth promotion by *E. meliloti* was lost in *nsp1* mutants under N-limiting conditions **(Figure 6A)**. Similarly, *atnlp4* and *atnlp9* mutant plants did not exhibit increased growth in the presence of *E. meliloti*. However, *atnlp4* mutant plants showed altered plant growth under all conditions evaluated. Conversely, *atnlp8* mutant plants did not affect the promotion of plant growth by *E. meliloti* under N-limiting conditions **(Figure 6A)**.

**Figure 6:**
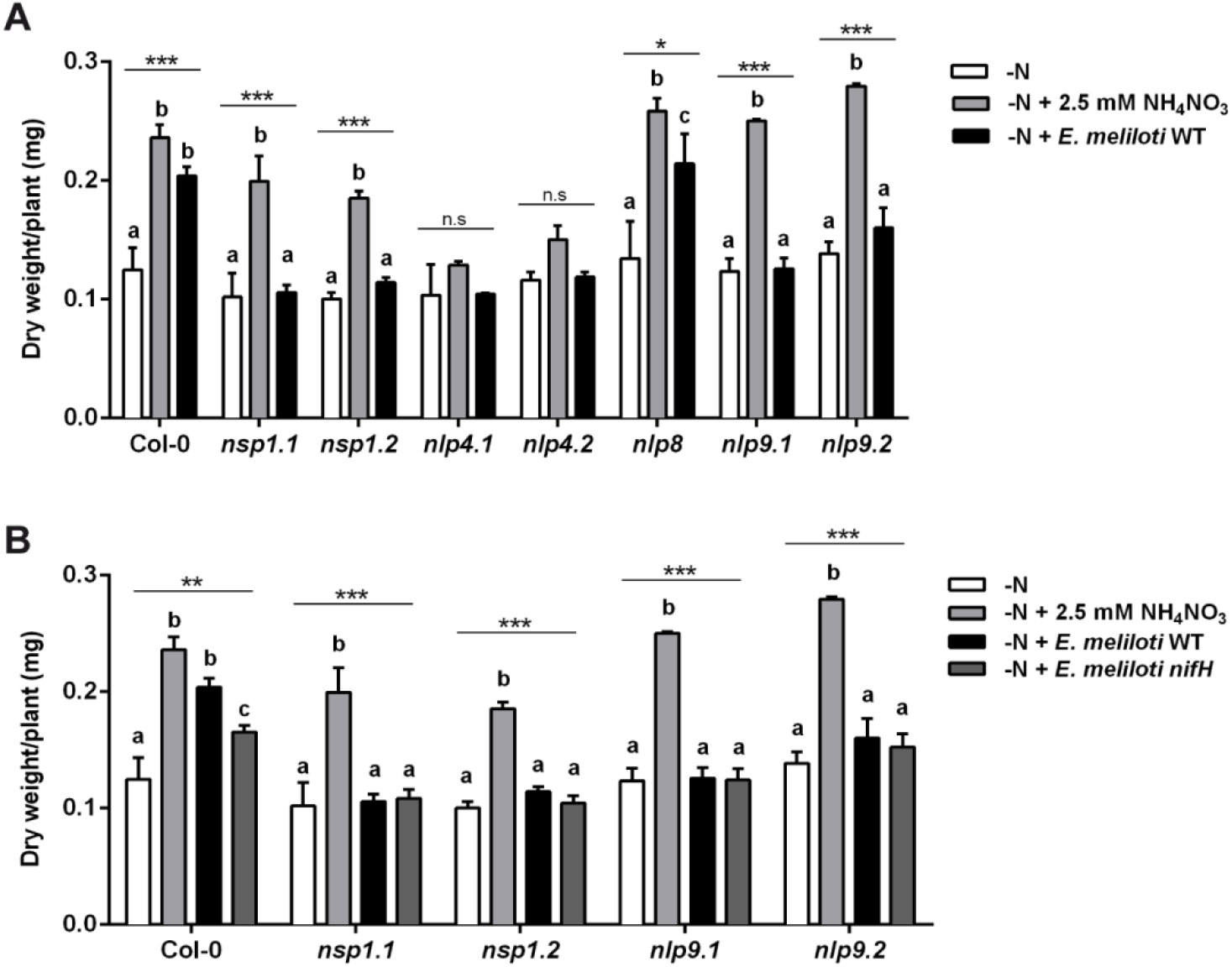
Putative transcription factors of *A. thaliana* are necessary for effective interaction with *E. meliloti*. Plants were grown and treated, as indicated previously. **(A)** Dry weight was measured in wild type, *nsp, nlp4, nlp8*, and *nlp9* mutant lines under limiting N conditions, limiting N conditions supplemented with 2.5 mM NH_4_NO_3_, and limiting N conditions inoculated with *E. meliloti* WT. **(B)** Dry weight was measured in wild type, *nspl.1, nspl.2, nlp9.1*, and *nlp9.2* mutant lines under limiting N conditions, limiting N conditions supplemented with 2.5 mM NH_4_NO_3_, and limiting N conditions inoculated with *E. meliloti* WT or *E. meliloti nifH*. Values plotted correspond to the mean of four independent biological replicates ± standard error. Results were subjected to two-way analysis of variance (ANOVA) and Tukey’s multiple comparison test. The letters indicate the means that differed significantly in each genotype, asterisks indicate p-value (*p < 0.05; ** p < 0.01; ***p<0.001; n.s, not significant).

These results suggest *AtNSP1* and *AtNLP9* are specifically required for a functional interaction between *A. thaliana* and *E. meliloti* for enhanced growth under N-limiting conditions.

Finally, to evaluate whether *AtNSP1* and *AtNLP9* are necessary for interaction with the bacteria or are required in the N-fixing process *nsp1.1, nsp1.2, nlp9.1*, and *nlp9.2* mutant plants were inoculated with WT, and *nifH E. meliloti* strains and the effect on biomass was determined. As shown in **Figure 6B**, the increased biomass retention that was observed in *nifH* (**Figure 2A)** was lost in these mutant plants.

These results suggest that *AtNSP1* and *AtNLP9* are required for effective *A. thaliana –E. meliloti* interaction under N-deficiency.

## DISCUSSION

In this study, we present biochemical, cell biological, physiological, and genetic evidence indicating that *E. meliloti* interacts with *A. thaliana* to promote plant growth under conditions of severe N-deficiency. This promotion of growth under N-deficiency is mediated in part by bacterial N-fixation. Our results demonstrate the rhizosphere of *A. thaliana* roots can be colonized by N-fixing bacteria. *A. thaliana* homologs of key regulatory genes involved in rhizobacteria-legume interactions such as *NLP* and *NSP* genes were required for growth promotion mediated by *E. meliloti*. Interestingly Nod factors genes were not required for this process. This bacterial-plant interaction is less efficient than the one found in legumes and provides limited N-nutrition and growth. However, it is very important because it allows the plant to complete its life cycle under severe N-deficiency.

Arabidopsis plant-growth enhancement in the absence of N specifically requires a metabolically active *E. meliloti* and is not caused by the mere presence of neutral or plant growth-promoting bacteria or other N-fixing bacteria. Both WT and *nifH* mutant bacterial strains increased lateral root density as compared with mock-inoculated plants indicating that additional bacterial growth-promoting pathways exist (Galleguillos et al., 2000). It is known that rhizobacteria, such as *E. meliloti*, have multiple mechanisms to promote plant growth. Previous studies have described a growth-promotion pathway in *E. meliloti*, which is independent of N-fixation (Chi et al., 2010) and can produce a signal molecule promoting growth in different plant species (Matiru and Dakora, 2005). However, it is important to note that these experiments were performed in the presence of N, which can inhibit biological N-fixation. Our assays were performed in the absence of N. Therefore, this general plant growth-promoting activity was not evaluated. Instead and most importantly, our study evaluated growth promotion due to biological N-fixation.

*A. thaliana* can functionally interact with *E. meliloti* to enhance plant growth under N-limiting conditions via N-fixation. Previous reports show plant growth-promoting effect on lateral-root development in *A. thaliana* inoculated with the Phyllobacterium strain STM196, but this effect was independent of the concentration of N in the media. However, acetylene reduction activity was evaluated in the presence of N in the media and, therefore, did not detect activity (Mantelin et al., 2006). More recent work analyzed the interaction between *A. thaliana* and *Mesorhizobium loti* (Poitout et al., 2017). They found an increase in shoot biomass production and transient inhibition of primary-root growth. Nevertheless, they did not perform the acetylene reduction assay or isotopic dilution experiments and only speculated that N-fixation is unlikely under their conditions (Poitout et al., 2017). We show *E. meliloti* WT can fix N when interacting with *A. thaliana* in the absence of N. While this interaction is less intimate than previous systems described in legumes or non-legume species, our results indicate it is sufficient to provide N for plant survival.

Even though there is no endophytic relationship between the plant and the bacteria, *E. meliloti* promotes growth and N nutrition of *A. thaliana* in the absence of a mineral N source. While we do not know the mechanism that could ensure an anoxic environment for biological N-fixation, several mechanisms that allow bacteria to isolate the nitrogenase complex from oxygen have been described (Ott et al., 2005; Bobik et al., 2006; Gupta et al., 2011). For instance, mucilaginous plant structures could allow N-fixation under aerobic conditions (Van Deynze et al., 2018). In addition to carbohydrates and amino acids, the mucilage secreted by the plant calyptra also contains specific chemoattractant components for beneficial microorganisms. In *A. thaliana*, the presence of arabinogalactans has been associated to the colonization of Rhizobium sp. in the root (Vicre et al., 2005). The formation of a biofilm by the bacteria has also been described (Wang et al., 2017). Interestingly, the catabolism of oxalate and arginine deaminase are activated to obtain energy under microaerobic conditions (Paungfoo-Lonhienne et al., 2016), suggesting that bacteria could create microaerobic conditions suitable for biological N-fixation. Moreover, host leghemoglobin proteins bind free oxygen to create a microoxic environment in N-fixing cells, protecting the nitrogenase enzyme from inactivation by oxygen (Kim et al., 2015).

Under our experimental conditions, *E. meliloti* colonizes the root surface of *A. thaliana* as rhizosphere bacteria. Although we cannot rule out location in the apoplast, our electron microscopy analysis did not show bacteria in intercellular spaces. Also, the analysis of distal and proximal agar samples suggests that the *NifH* gene is activated when bacteria are in the proximity of the root. Transcription of *NifH* does not necessarily mean that it is translated into a functional protein. However, this evidence taken together with our other results suggests that the interaction occurs when the plant and the bacteria are close.

Previous studies reported a diversity of bacterial communities associated with Arabidopsis, including many species with potential N-fixation capabilities. In some studies, intercellular root colonization of *A. thaliana* has been shown; in other cases, diazotrophic bacteria could ectopically colonize leaves and roots (Gough et al., 1997; Stone et al., 2001; Bulgarelli et al., 2012; Lundberg et al., 2012; Bai et al., 2015). Moreover, potentially N-fixing bacteria have been found in the microbiome of non-legume plants (e.g., Acidovorax strains and *Paraburkholderia kururiensis*) (Levy et al., 2018). Even though these results do not prove these bacteria are competent in N-fixation as part of the Arabidopsis microbiota, they showed that diazotrophic bacteria can colonize *A. thaliana* and are consistent with our results.

Although some studies have shown that Rhizobiales are consistently found in high relative abundances and enriched in the root and leaf communities of phylogenetically diverse plant hosts, lack of nodulation and N-fixation capability in bacterial isolates from roots have also been noted (Garrido-Oter et al., 2018). It is possible that an ancestral mechanism of interaction with both non-leguminous and leguminous hosts enables successful root colonization. The majority of tested strains elicited robust root growth promotion in *A. thaliana* and consistently rescued phosphate starvation-induced root-growth inhibition and retained root-growth promotion under nitrogen starvation, suggesting that unlike in nodule symbiosis part of the Arabidopsis microbiota operate both under nitrogen-sufficient and - deficient conditions (Garrido-Oter et al., 2018).

Based on previous and results of this work, we propose a gradient of plant-rhizobacteria interactions, of which ours would represent an early evolutionary step for biological N-fixation. The more intimate relationships between plants and rhizobacteria are exemplified by specialized organ structures such as nodules found in legumes. This has also been observed in other non-legume plants. Commercial crops such as wheat and sugarcane and others such as *Setaria viridis* can assimilate a significant part of their N requirements through BNF when exposed to N-limiting conditions (Boddey et al., 1995; Iniguez et al., 2004; Pankievicz et al., 2015). A recent study in maize hypothesized that isolated indigenous landraces of maize, grown with little or no fertilizer, might have evolved strategies to improve plant performance under low-nitrogen nutrient conditions (Van Deynze et al., 2018). They showed these landraces, characterized by the extensive development of aerial roots, secrete carbohydrate-rich mucilage that can harbor diazotrophic microbiota. Under these conditions, they found that 29%–82% of the plant N is derived from atmospheric nitrogen (Van Deynze et al., 2018).

N-fixation in legume species is regulated by molecular pathways that control when and how the symbiotic association is established with rhizobia. Plant genes involved in this interaction have been characterized in legumes (Oldroyd, 2013). However, in Brassicales such as Arabidopsis, some of the symbiosis-specific genes have been lost (Delaux et al., 2014). A set of conserved genes present in Arabidopsis, including Nodulation-Signaling Pathway (NSPs) (Delaux et al., 2014) and NIN-like protein (NLPs) genes, have been evaluated (Schauser et al., 2005). *AtNSP1* function is required under N-limiting conditions when *E. meliloti* is present. Potential orthologs of NSP1 and NSP2 can be found in many higher plant species, including rice (*Oryza sativa*) (Liu et al., 2011). On the other hand, genetic studies in *Lotus japonicus* and pea have identified *NIN* as a core symbiotic gene required for establishing symbiosis between legumes and N-fixing bacteria (Schauser et al., 2005). *A. thaliana* has 9 NIN Like Proteins (NLPs) (Ott et al., 2005), of which *AtNLP6* and *AtNLP7* play a key role in nitrate-signaling by regulating nitrate-inducible gene expression (Marchive et al., 2013; Guan et al., 2017). This work suggests *AtNSP1* and *AtNLP9* are necessary for *A. thaliana* – *E. meliloti* interaction and are also required in the N-fixing process, in a Nod factor-independent manner.

We propose *E. meliloti* and *A. thaliana* can interact as a salvage mechanism, which allows the plant to procure N under extreme N-deficiency by recruiting N-fixing bacteria to the rhizoplane and thus escape adverse conditions. Under these circumstances, N supply mediated by N-fixation is critical for survival, allowing the plant to complete its life cycle.

This work, along with previously published studies, suggests an expanding role for beneficial plant-bacteria interactions for N-nutrient acquisition. It can contribute to catalyzing further efforts to understand the underlying mechanisms and ecological consequences of this phenomenon. In this regard, *A. thaliana*-*E. meliloti* interaction represents an excellent model system to address non-legume plant mechanisms to promote interactions with N-fixing bacteria, which may eventually lead to the development of biotechnologies for more sustainable strategies for plant N nutrition.

## METHODS

### Plants and bacteria

*Arabidopsis thaliana* Columbia (Col-0) ecotype was used in all experiments. T-DNA insertional mutant plants for *NSP* and *NLP* genes were obtained from the Arabidopsis Biological Resource Center (https://abrc.osu.edu/). Homozygous lines were selected using PCR, the T-DNA primer LBa1 (5’-TGGTTCACGTAGTGGGCCATCG-3’), and the gene-specific primers described in **Table 1**.

**Table 1:**
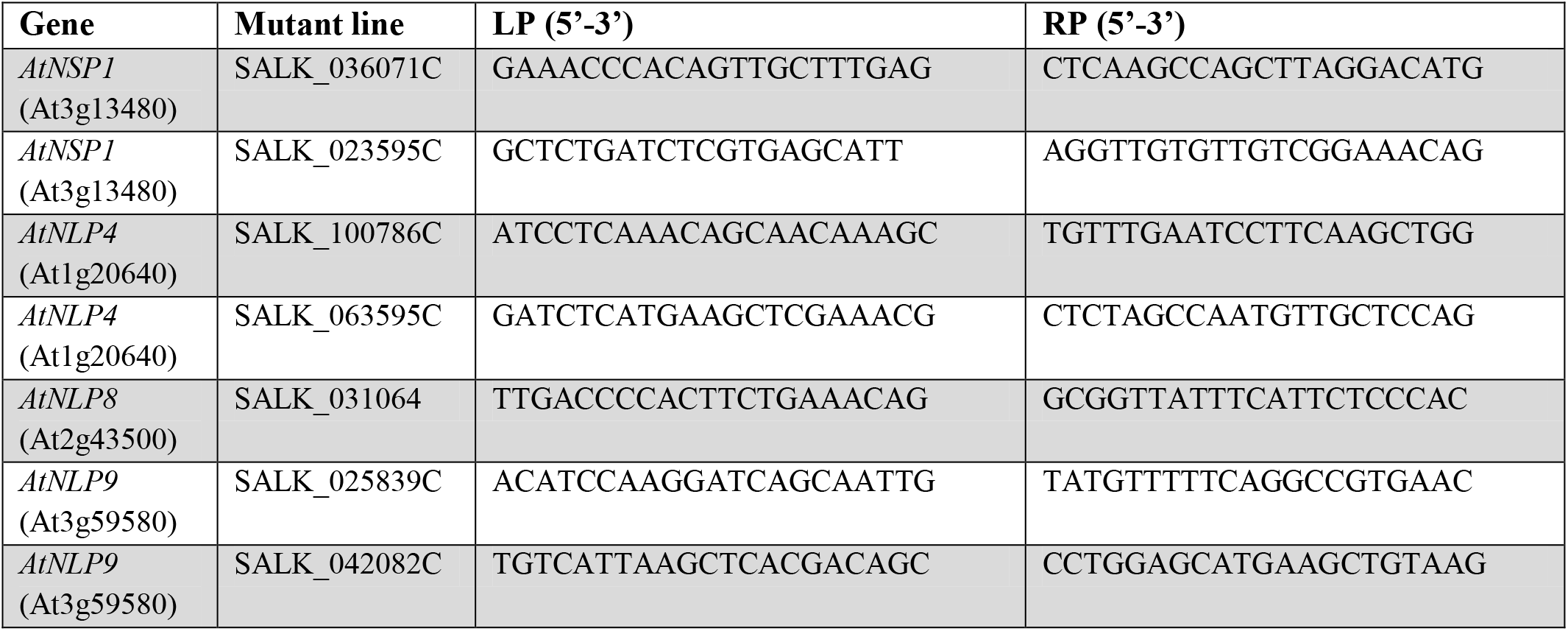
Primers used for the genotyping of *A. thaliana* mutant lines.

For evaluation of *E. meliloti* mutant strains, Alfalfa plants (*Medicago sativa* cv. Vernal) were used.

Bacterial strains utilized were *Rhizobium etli* CFN42 (Poupot et al., 1995), *Cupriavidus taiwanensis* LMG19424 (Marchetti et al., 2011), *Paraburkholderia xenovorans* LB400 (Sawana et al., 2014), *Paraburkholderia vietnamiensis* G4 (Sawana et al., 2014), *Paraburkholderia phytofirmans* PsJN (Zuniga et al., 2013; Sawana et al., 2014), *Cupriavidus pinatubonensis* JMP134 (Ledger et al., 2012), *Ensifer meliloti* RMP110 WT (Yuan et al., 2006), and its derivatives *nifH, nodA*, P_*Ni*fH_::*GFP* and P_*lac*Iq_::*GFP* (this study).

### E. meliloti gene inactivation and construction of plasmid derivatives expressing GFP reporter gene

*E. meliloti Nif*H and *NodA* genes were inactivated according to the procedure described by Louie et al. (2002). Each gene was disrupted via homologous integration of a suicidal plasmid that carried the internal fragment of the respective gene. A 295-bp internal fragment of the *NifH* gene and a 568-bp internal fragment of the *Nod*A gene were amplified from *E. meliloti* DNA by using the specific primers described in **Table 2**. The PCR products were cloned into pCR2.1-TOPO (Invitrogen, Carlsbad, CA, USA). Final plasmids DNA (5-10μg) were electroporated into electrocompetent *E. meliloti* cells and neomycin (200 μg/mL) selection was used to identify the transformed bacteria. PCR amplification and direct sequencing were carried out to verify plasmid integration and gene interruption in *nifH* and *nodA* mutant *E. meliloti* strains.

**Table 2:**
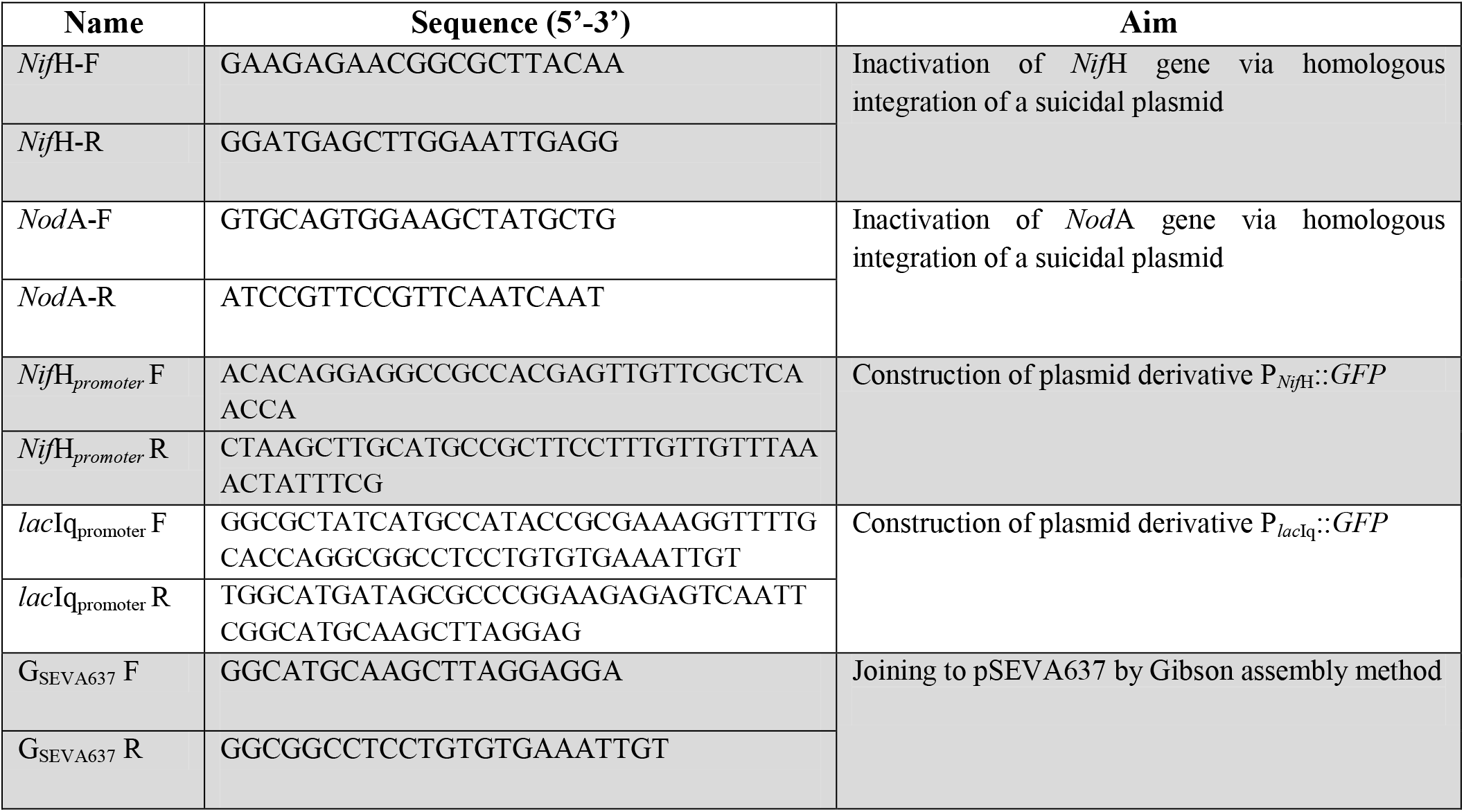
Primers used for the inactivation of *E. meliloti* genes and the construction of plasmids expressing the GFP reporter gene.

For the construction of plasmid derivatives expressing the green fluorescent protein (GFP) reporter, pSEVA plasmids belonging to Standard European Vector Architecture were utilized (Silva-Rocha et al., 2013). PCR products comprising *NifH* gene promoter sequence from *E. meliloti* and *lacIq* promoter sequence from pSEVA614 plasmid (GenBank: JX560386.2)(Silva-Rocha et al., 2013), were obtained by using the specific primers described in **Table 2** and joined to pSEVA637 low copy number plasmid (origin of replication pBBR1, Gm selection marker, and GFP) (Silva-Rocha et al., 2013) in a one-step isothermal reaction by Gibson assembly method (Gibson et al., 2009). Inducible reporter plasmid pSEVA-P_*NifH*_-GFP and constitutive promoter plasmid pSEVA-P_*lac*Iq_-GFP were generated. These recombinant plasmids were electroporated into *E. meliloti*, and gentamicin (10 μg/mL) selection was used to identify the transformed bacteria *E. meliloti* P_*NifH*_::*GFP* and P_*lac*Iq_::*GFP*.

### Preparation of bacterial samples for inoculation

All bacteria were routinely grown with 869 medium diluted 1/10 (1 liter of non-diluted medium contains: 1g tryptone, 0.5g yeast extract, 0.5g NaCl, 0.1g D-glucose, and 0.0345g CaCl_2_xH_2_O) (Mergeay et al., 1985) in an orbital shaker (200 rpm) for 48h-72h at 28°C to an optical density (OD_600_nm) value of 0.4. Then, cells from 20 mL of culture were harvested by centrifugation, washed three times, and resuspended in 5 mL sterile water. The final suspension for each strain was homogeneously spread on 20 mL of 0.8% total agar plates containing Murashige and Skoog (MS) basal salt mixture without nitrogen (M531; Phytotechnology Laboratories) as indicated for each experiment. For the analysis with dead bacteria, before inoculation, bacterial cells were incubated at 100°C for 10 min and then resuspended in 5 mL sterile water and diluted to 20 mL of total plate volume.

### Plant growth promotion assays

*A. thaliana* seedlings grown on vertical plates with 5mM KNO_3_ were transferred at the end of the seventh day to MS basal salt mixture without N, supplemented with 2.5mM NH_4_NO_3_ or inoculated with bacteria. After one week of incubation, plants were harvested and dried at 70°C for two days.

Biomass was measured as the dry weight of thirty seedlings (which corresponds to one biological replicate) and normalized by the number of plants. Every experiment was done with at least three independent biological replicates for each condition. All experiments were performed under long days (16/8 h light/dark cycles) at 22°C in Percival incubators model CU36L5.

For the evaluation of *E. meliloti nifH* and *nodA* mutant strains, Alfalfa seeds (*Medicago sativa* cv. Vernal) were superficially sterilized with 50% sodium hypochlorite solution and sown in MS media without N and inoculated with *E. meliloti* WT, *nifH*, or *nodA* mutant strains. Nodule formation and phenotype were evaluated four weeks after inoculation.

### ^15^N dilution and acetylene reduction assays

For ^15^N dilution assay, Arabidopsis seedlings were grown vertically in MS medium without N and supplemented with 5 mM isotopically labeled KNO_3_ (5% of ^15^N). After seven days, plants were washed with 0.01 mM CaSO_4_ and transferred to MS media without N or supplemented with 2.5 mM NH_4_NO_3_ as the only N source, or without N inoculated with WT or *nifH* mutant *E. meliloti* strains. Whole plants were harvested seven days after the transfer and dried for three days at 70°C. Plant samples were sent to the mass spectrometry facility at Elemental Analysis Service of the Laboratory of Biogeochemistry and Applied Stable Isotopes (LABASI) (P. Universidad Católica de Chile) to determine the abundance of elemental ^15^N isotopes on an Isotope Ratio Mass Spectrometer, IRMS, Thermo Delta Advantage coupled to an EA2000 Flash Elemental Analyzer.

The normalization procedure described in Putz et al. (2011) was used to determine the atom percent excess (APE), whereby the enrichment of plants post-labeling was calculated by subtracting the mean atom% value of the control plants from the atom% of the labeled plants, yielding atom% excess values (APE).

For the acetylene reduction assay (ARA), Arabidopsis seedlings were grown vertically in MS medium without N and supplemented with 5 mM KNO_3_, as described previously. Seven days after germination, plants were transferred to 250 mL hermetically-sealed flasks (with a rubber stopper) containing MS medium without N and inoculated with WT or *nif*H mutant *E. meliloti* strains. We used MS medium without N or supplemented with 2.5 mM NH_4_NO_3_ as controls. After two weeks, flasks were sent to the Laboratory of Biogeochemistry and Applied Stable Isotopes (LABASI) (P. Universidad Católica de Chile) to perform ARA, according to the protocol described by Hardy et al. (1968). The treated samples and the controls were injected with acetylene, generating a 10% v/v mixture of acetylene/air. No acetylene was added to one flask in each treatment to control the possible production of ethylene by processes unrelated to nitrogenase activity. After 24 hours, an air sample was taken from the flasks, and a gas chromatograph was used to determine the concentration of ethylene generated. After testing, the samples were dried at 60 °C for at least 48 h, to obtaining their dry weight. The acetylene reduction rates were obtained by estimating the slope of the ethylene concentration curve as a function of the time of incubation using the dry weight of the incubated sample as a reference. The ethylene nanomolar produced per hour of incubation and gram of dry weight (nmol ethylene h-1 g-1 dry weight) was estimated. Controls were subtracted from the concentration detected in the treatments (Pérez et al., 2017a; Pérez et al., 2017b).

### Microscopy analysis

Microscopy analysis was performed using five different experimental approaches. (**A**) For optical microscopy, Arabidopsis plants incubated with bacterial cells as described above were fixed in 2.5% glutaraldehyde, 0.268 M sodium cacodylate buffer at pH 7.2. A selection of root tips and axillary areas were obtained with a microtome to achieve semi-thin longitudinal cuts. These cuts were subsequently stained with toluidine blue. This procedure was carried out by the advanced microscopy unit (UMA) of the Pontificia Universidad Católica de Chile. The slices were analyzed in a Nikon ECLIPSE Ni optical microscope. (**B**) For Transmission Electron Microscopy (TEM) analysis, the same root tips and axillary areas fixed in glutaraldehyde were used to obtain thin longitudinal cuts, mounted on a grid, stained with uranyl acetate by UMA, and later analyzed using a Philips Tecnai 12 transmission electron microscope (Biotwin). (**C**) For Scanning Electron Microscopy (SEM) analysis, plant roots incubated with bacteria were fixed in 2.5% glutaraldehyde, dehydrated, subjected to critical point drying with acetone/CO_2_ and then shaded with gold by UMA. The observation was made in a Hitachi TM3000 scanning microscope at 15 kV. (**D**) For confocal laser scanning microscopy, to visualize bacterial GFP expression, epidermal root cells were stained with propidium iodide (50μg/mL). Samples were analyzed using an Olympus LSM Fluoview 1000 system confocal microscope, multi Argon gaseous laser 40 mW 488 nm and 543 nm to observe GFP (green) and propidium iodide (red), respectively. (**E**) To analyze bacterial *Nif*H expression, Arabidopsis plants were incubated with bacterial cells carrying a *pNifH::GFP* reporter gene in the presence and absence of nitrogen in the medium. An agar sample from an area adjacent to the root and another sample 5 cm away were mounted on a slide and analyzed by confocal laser scanning microscopy using the same laser parameters indicated above. GFP fluorescence was normalized, first, by extracting possible autofluorescence from the agar, using a plate without bacteria. Then, for each agar sample, 13 slices were taken on the Z-axis. Images were then integrated using the ImageJ FIJI program (Schneider et al., 2012). A fluorescence threshold was defined; signals over that value were quantified and normalized by bacterial area. 100 of these normalized measurements were randomly selected for each experimental condition, and their frequency, density, and relative fluorescence were determined.

### RNA isolation and gene expression analysis

RNA extraction was performed with the Ambion PureLink_TM_ RNA Mini Kit. cDNA synthesis was carried out using the Improm-II reverse transcriptase according to Manufacturer’s instructions (Promega). Gene expression analysis was carried out using the Brilliant SYBR Green QPCR Reagents on a Stratagene MX3000P qPCR system according to the instructions of the manufacturer and the primers described in **Table 3**. RNA levels were normalized to *ADAPTOR PROTEIN-4 MU-ADAPTIN* (At4g24550) gene expression.

**Table 3:**
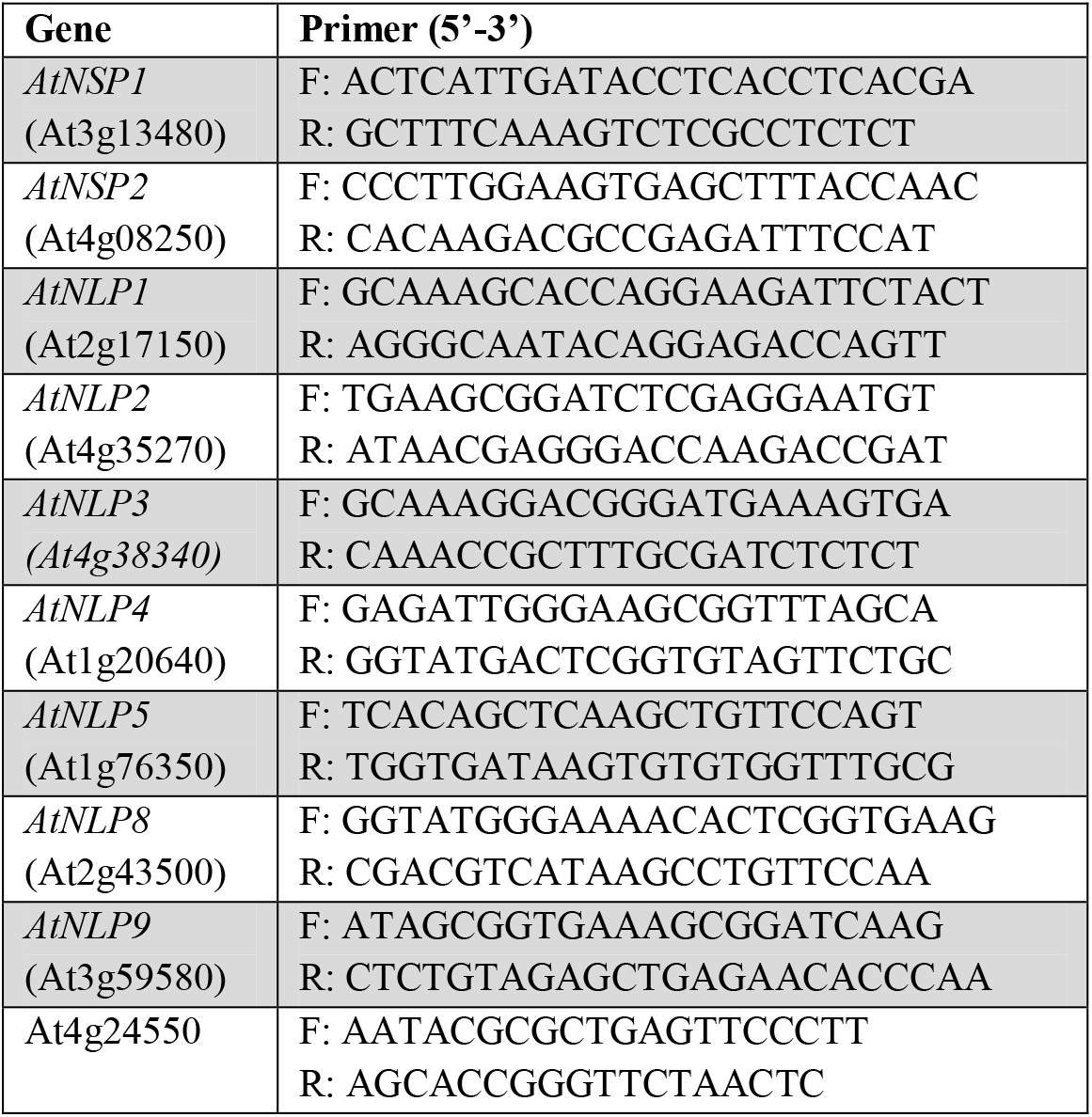
Primers used for the qRT-PCR of *A. thaliana* genes.

### Statistical Analysis

All data presented in this study correspond to the mean ± SE of at least three biological replicates for each sample. Results were subjected to a one-way or two-way analysis of variance (ANOVA) and Tukey’s multiple comparisons test.

## FUNDING

This work was supported by, ANID/FONDAP/15090007, Instituto Milenio iBio – Iniciativa Científica Milenio MINECON, Fondo Nacional de Desarrollo Científico y Tecnológico (FONDECYT) 1180759.

## AUTHOR CONTRIBUTIONS

GA, T.K. and M.P.M. performed experiments. D.G. contributed complementary experimental work. G.A., T.K., M.P.M, B.G. and R.A.G designed experiments. R.A.G. supervised the study and analyzed results. A.Z. and B.G. collaborated in experimental design, troubleshooting and generating GFP-lines of bacteria. G.A., T.K. and R.G wrote the manuscript. The authors declare no conflict of interest.

## Notes

### Competing Interest Statement

The authors have declared no competing interest.

