## Supplemental Figure 1 for "*Arabidopsis thaliana* interaction with *Ensifer meliloti* can support plant growth under N-deficiency"

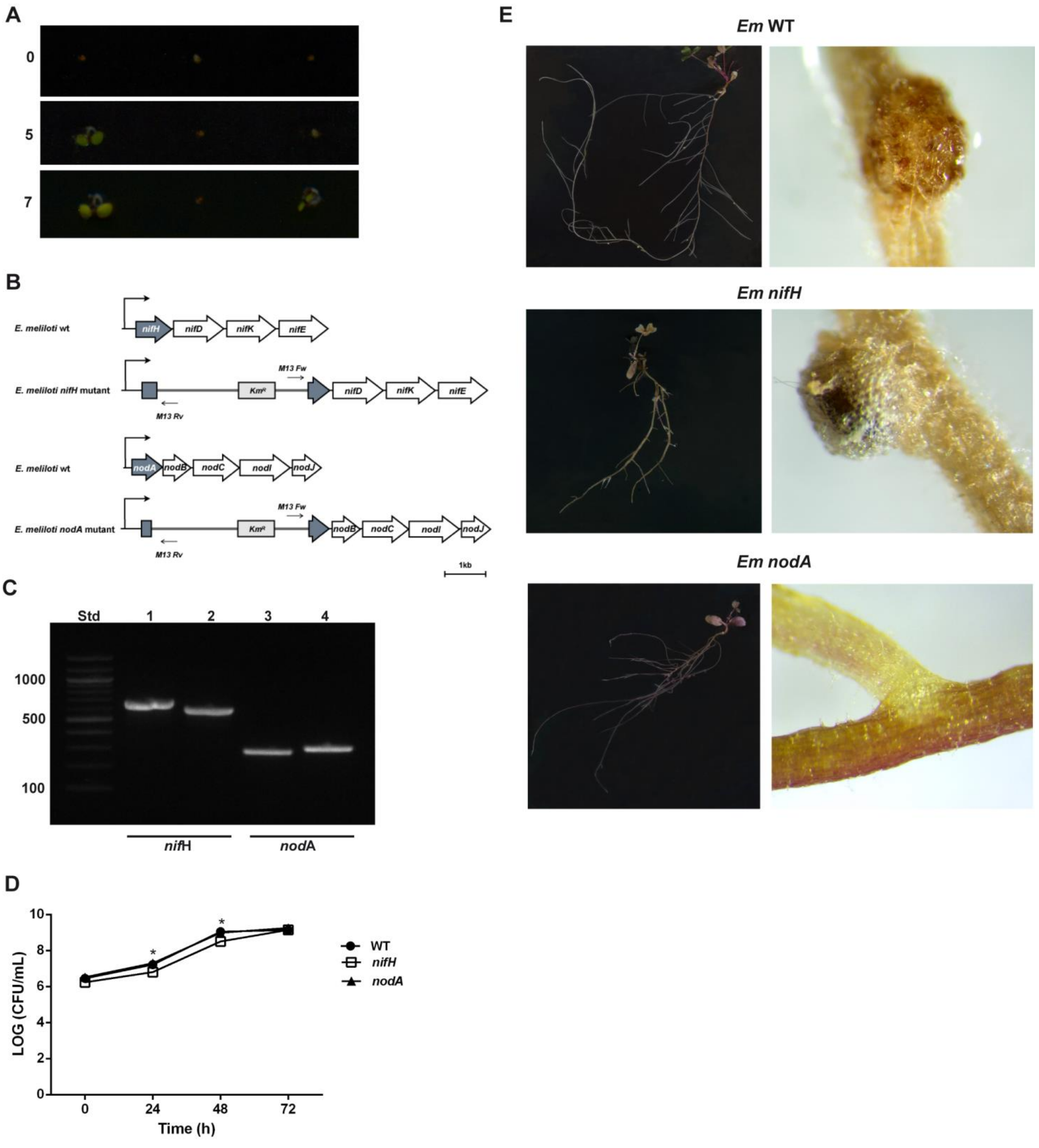


**Supplemental Figure 1: Phenotypic analysis of plants and genetic confirmation of mutant bacteria.**

**(A)** Germination of *A. thaliana* seeds obtained from plants grown in the absence of N, inoculated with *E. meliloti: A. thaliana* seeds, obtained from plants grown for eight weeks in MS media without N and inoculated with *E. meliloti*, were superficially sterilized with 50% sodium hypochlorite solution and sowed in MS ½. Germination of seeds was evaluated at 0, 5, and 7 days after sowing. **(B)** Scheme of the gene disruptions carried out for the mutant *E. meliloti* bacteria *nifH* and *nodA*. By disrupting the structural component of nitrogenase, NifH nitrogenase Fe protein (*NifH* gene), we aimed to disrupt the functionality of the entire cluster. Similarly, we disrupted the acyltransferase NodA, (*NodA* gene), the first gene in the Nod cluster, to impair the synthesis of Nod signal molecules in the strain.

**(C)** Evaluation of *E. meliloti nifH* and *nodA* mutant strains by PCR. Std: 100 bp standard. Lanes 1 and 2: PCR product for *nifH* amplification in mutant bacteria with *nifH*Fw & M13Rv and M13Fw & *NifH*Rv primers, respectively. Lanes 3 and 4: PCR product for *nodA* amplification in mutant bacteria with *nodA*Fw & M13Rv and M13Fw & *NodA*Rv primers, respectively. (*nifH*Fw: 5'-GTCCACGACCTCCCAAAATA-3'; *NifH*Rv: 5'-ATCTGCTCGTCGCTCTTCAT-3' ; M13Fw: 5'-GTAAAACGACGGCCAG-3'; M13Rv: 5'-CAGGAAACAGCTATGAC-3'; *nodA*Fw: 5'-GTGCAGTGGAAGCTATGCTG-3'; *NodA*Rv: 5'-ATCCGTTCCGTTCAATCAAT-3').

**(D)** To evaluate the viability of *E. meliloti nifH* and *nodA* mutant strains, 1% of bacterial culture in 100 mL of 869 media (200 ug/mL streptomycin) was incubated in an orbital shaker (200 rpm) at 28°C. Every 24 hours, a sample of the culture was taken, serially diluted, and plated in 869 with the corresponding antibiotic. After 48 hours, the forming colony units (CFU) were quantified. Values plotted correspond to the mean of three independent biological replicates ± standard error. Results were subjected to one-way analysis of variance (ANOVA) (*$p < 0.05$).

**(E)** Alfalfa plants (*Medicago sativa* cv. Vernal) were used to evaluate *E. meliloti nifH* and *nodA* mutant strains. For this, alfalfa seeds were superficially sterilized with 50% sodium hypochlorite solution and sowed in MS media without N and inoculated with *E. meliloti* WT, *nifH*, or *nodA* mutant strains. Nodule formation and phenotype were evaluated four weeks after inoculation.
