## Supplemental Figure 2 for "*Arabidopsis thaliana* interaction with *Ensifer meliloti* can support plant growth under N-deficiency"

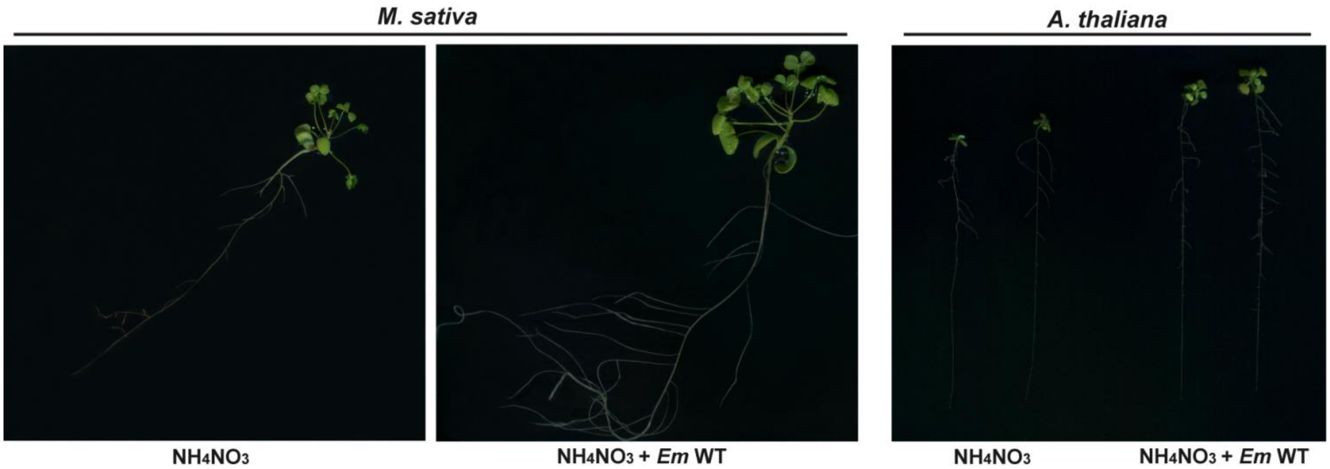


**Supplemental Figure 2: Evaluation of *E. meliloti* plant growth-promoting activity, independent of N-fixation.**
Alfalfa (*Medicago sativa* cv. Vernal) and *A. thaliana* plants were sowed in MS media without N supplemented with 2.5mM $NH_4NO_3$ and MS media without N inoculated with *E. meliloti* WT strain. Plants were kept in these conditions for 3 and 2 weeks for Alfalfa and Arabidopsis, respectively.
