## Supplemental Figure 3 for "*Arabidopsis thaliana* interaction with *Ensifer meliloti* can support plant growth under N-deficiency"

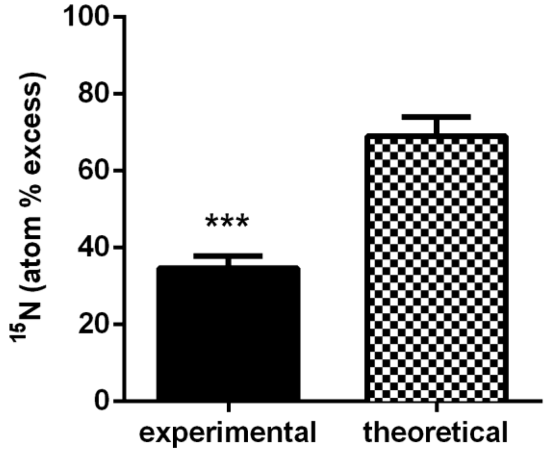


**Supplemental Figure 3: Theoretical estimate of total N contribution of inoculated *E. meliloti.***
To evaluate the contribution of *E. meliloti* biomass in $^{15}$N dilution assay, bacterial cultures ($OD_{600}$ 0.4) were harvested by centrifugation and washed three times with sterile water. The pellets obtained were dried for three days at 70°C, and the total N content was analyzed by the Elemental Analysis Service of the Laboratory of Biogeochemistry and Applied Stable Isotopes (LABASI) (P. Universidad Católica de Chile). Using the total N (mg) of *E. meliloti* inoculum and N content of *A. thaliana*, we estimated a theoretical $^{15}$N atom percent excess. For this calculation, we considered *A. thaliana* absorbing 100% of the N contained in *E. meliloti* biomass and a $^{15}$N natural abundance of 0.3663%, using the $^{15}$N atom percent excess determined in non-inoculated plants as a reference for undiluted conditions (Figure 2c).
Results were subjected to t-test stadistical analysis. (***$p < 0.001$).
