## Supplemental Figure 4 for "*Arabidopsis thaliana* interaction with *Ensifer meliloti* can support plant growth under N-deficiency"

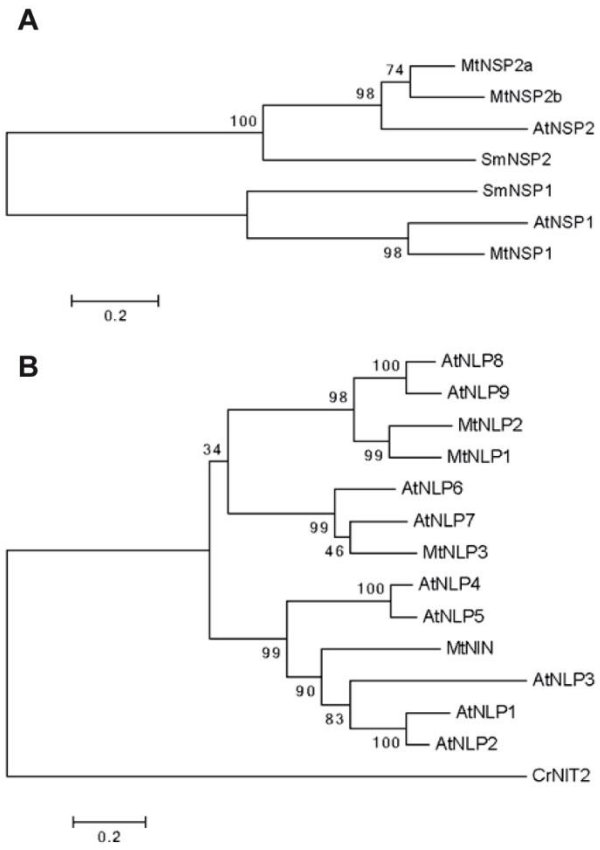


**Supplemental Figure 4: Evolutionary relationship between *A. thaliana* and *M. truncatula* NSPs and NLPs.**

Phylogenetic tree of NSPs **(A)** and NLPs **(B)** proteins. Sequences from *A. thaliana*, *M. truncatula*, *Chlamydomonas reinhardtii*, and *Selaginella moellendorffii* proteins were aligned using MAFFT software. The phylogenetic trees were computed with MEGA6 software using maximum likelihood and bootstrap with 5000 replicates.
